# Enhancer hijacking determines intra- and extrachromosomal circular *MYCN* amplicon architecture in neuroblastoma

**DOI:** 10.1101/2019.12.20.875807

**Authors:** Konstantin Helmsauer, Maria Valieva, Salaheddine Ali, Rocio Chamorro Gonzalez, Robert Schöpflin, Claudia Röefzaad, Yi Bei, Heathcliff Dorado Garcia, Elias Rodriguez-Fos, Montserrat Puiggròs, Katharina Kasack, Kerstin Haase, Luis P. Kuschel, Philipp Euskirchen, Verena Heinrich, Michael Robson, Carolina Rosswog, Jörn Tödling, Annabell Szymansky, Falk Hertwig, Matthias Fischer, David Torrents, Angelika Eggert, Johannes H. Schulte, Stefan Mundlos, Anton G. Henssen, Richard P. Koche

## Abstract

*MYCN* amplification drives one in six cases of neuroblastoma. The supernumerary gene copies are commonly found on highly rearranged, extrachromosomal circular DNA. The exact amplicon structure has not been described thus far and the functional relevance of its rearrangements is unknown. Here, we analyzed the *MYCN* amplicon structure and its chromatin landscape. This revealed two distinct classes of amplicons which explain the regulatory requirements for *MYCN* overexpression. The first class always co-amplified a proximal enhancer driven by the noradrenergic core regulatory circuit (CRC). The second class of *MYCN* amplicons was characterized by high structural complexity, lacked key local enhancers, and instead contained distal chromosomal fragments, which harbored CRC-driven enhancers. Thus, ectopic enhancer hijacking can compensate for the loss of local gene regulatory elements and explains a large component of the structural diversity observed in *MYCN* amplification.

## Introduction

Oncogene amplification is a hallmark of cancer genomes. It leads to excessive proto-oncogene overexpression and is a key driver of oncogenesis. The supernumerary gene copies come in two forms, i. self-repeating arrays on a chromosome (homogeneously staining regions, HSR) and ii. many individual circular DNA molecules (extrachromosomal DNA, ecDNA, alias double minute chromosomes, dmin)^1^. EcDNA can arise during genome reshuffling events like chromothripsis and are subsequently amplified^2,3^. This partially explains why such circular DNAs can consist of several coding and non-coding distant parts of one or more chromosomes^4^. Over time, amplified DNA acquires additional internal rearrangements as well as coding mutations, which can confer adaptive advantages such as resistance to targeted therapy^5–7^. EcDNA re-integration into chromosomes can lead to intrachromosomal amplification as HSRs^8,9^ and act as a general driver of genome remodeling^10^. Our knowledge of the functional relevance of non-coding regions co-amplified on ecDNA, however, is currently limited.

*MYCN* amplification is a prototypical example of a cancer-driving amplification. The developmental transcription factor was identified as the most commonly amplified gene in a recent pediatric pan-cancer study^11^. Its most prominent role is in neuroblastoma, a pediatric malignancy of the sympathetic nervous system. *MYCN* amplification characterizes one in six cases and confers dismal prognosis^12^. In contrast to long-term survival of more than 80% for non-amplified cases, 5-year overall survival is as low as 32% for *MYCN*-amplified neuroblastoma^12^. In these cases, *MYCN* amplification is likely an early driver of neuroblastoma formation. Accordingly, *MYCN* overexpression is sufficient to induce neuroblastic tumor formation in mice^13,14^. Despite its central role in neuroblastoma biology, the epigenetic regulation of *MYCN* is incompletely understood.

Recently, studies have identified a core regulatory circuit (CRC) including half a dozen transcription factors that drive a subset of neuroblastoma with noradrenergic cell identity, including most *MYCN*-amplified cases^15–18^. The epigenetic landscape around *MYCN* is less well described. In part, this is due to the structural complexity of *MYCN* amplicons and difficulties in the interpretation of epigenomic data in the presence of copy number variation. Recent evidence has emerged suggesting that local enhancers may be required for proto-oncogene expression on amplicons^19^. Here, we sought out to identify key regulatory elements near *MYCN* in neuroblastoma by integrating short- and long-read genomic and epigenomic data from neuroblastoma cell lines and primary tumors. We investigated the activity of regulatory elements in the context of *MYCN* amplification and characterized the relationship between amplicon structure and epigenetic regulation.

## Results

### Local CRC-driven enhancers contribute to MYCN expression in neuroblastoma

In order to identify candidate regulatory elements near *MYCN*, we examined public H3K27ac chromatin immunoprecipitation and sequencing (ChIP-seq) and RNA sequencing (RNA-seq) data from 25 neuroblastoma cell lines^15^. ChIP-seq data for amplified genomic regions are characterized by a very low signal-to-noise ratio, which has complicated their interpretation in the past^16^. We therefore focused our analysis on 12 cell lines lacking *MYCN*-amplifications but expressing *MYCN* at different levels, allowing for the identification of *MYCN*-driving enhancers in neuroblastoma. Comparison of composite H3K27ac signals of *MYCN*-expressing vs. non-expressing cell lines identified at least 5 putative enhancer elements (e1-e5) that were exclusively present in the vicinity of *MYCN* in cells expressing *MYCN*, thus likely contributing to *MYCN* regulation (Fig. 1, Supplementary Fig. 1a). Consistent with differential RNA expression, a strong differential H3K27ac peak was identified spanning the *MYCN* promotor and gene body (MYCNp; Fig. 1). The identified enhancers were not active in developmental precursor cells such as embryonic stem cells, neuroectodermal cells or neural crest cells (Supplementary Fig. 1b), suggesting these enhancers were specific for later stages of sympathetic nervous system development or neuroblastoma. Transcription factor ChIP-seq in *MYCN*-expressing cells confirmed that four of the enhancers (e1, e2, e4, e5) were bound by each of three noradrenergic neuroblastoma core regulatory circuit factors (PHOX2B, HAND2, GATA3; Fig. 1b). All but enhancer e3 harbored binding motifs for the remaining members of the core regulatory circuit (ISL1, TBX2, ASCL1; Supplementary Fig. 1c) for which ChIP-seq data were unavailable. Additionally, all enhancers contained binding motifs for TEAD4, a transcription factor implicated in a positive feedback loop with MYCN in *MFCN*-amplified neuroblastoma^20^. Two of the enhancers (e1, e2) also harbored canonical E-boxes, suggesting binding of MYCN at its own enhancers (Supplementary Fig. 1c). Thus, a common set of CRC-driven enhancers is found specifically in *MYCN* expressing neuroblastoma cells, indicating that *MYCN* expression is regulated by the CRC.

**Figure 1.**
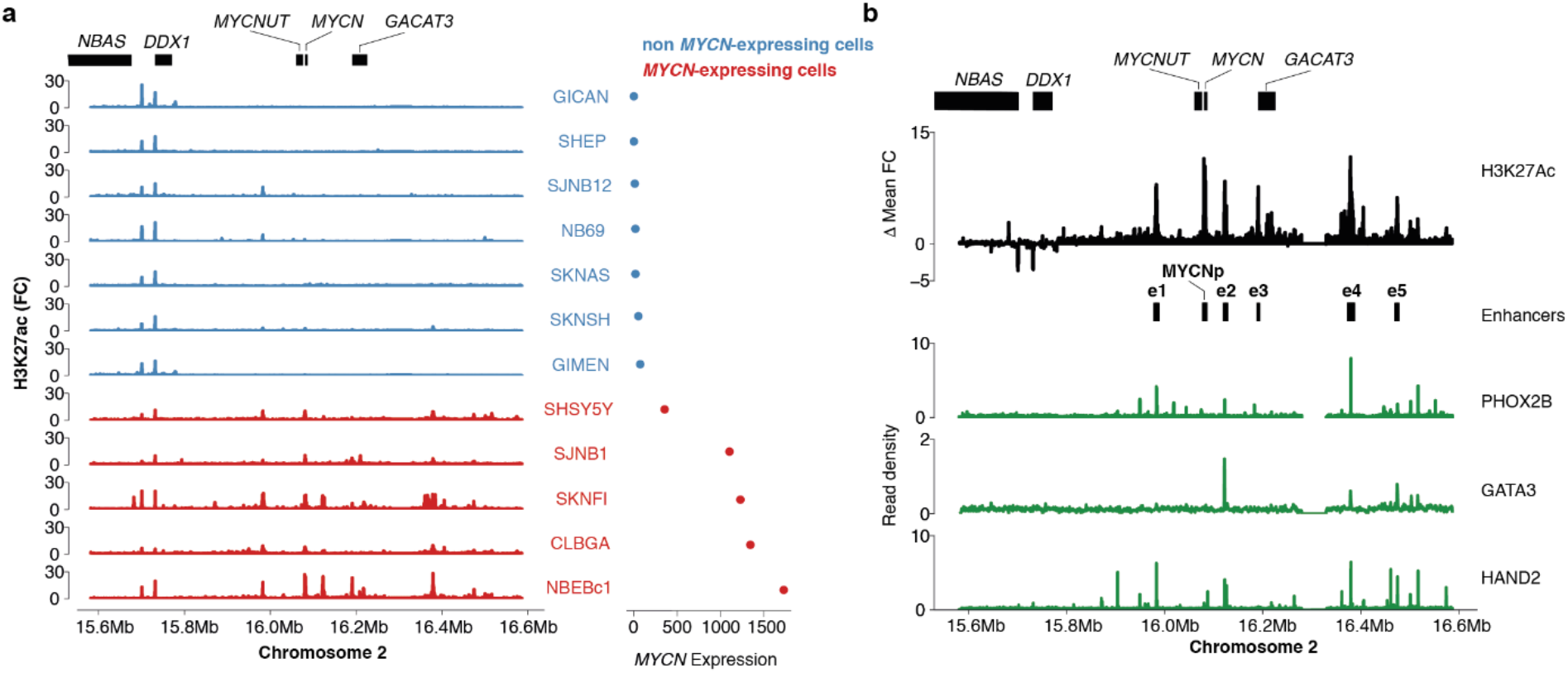
Five enhancers are specifically found in *MYCN*-expressing neuroblastoma cells. **a,** H3K27ac ChIP-seq fold change over input (left) and size-factor normalized *MYCN* expression as determined from RNA-seq for 12 non-MYCN-amplified neuroblastoma cell lines (*MYCN*-expressing, red; non *MYCN*-expressing, blue). **b,** Aggregated H3K27ac signal of *MYCN*-expressing compared to non-expressing cells (top; MYCNp, *MYCN* promotor; e1-e5, *MYCN*-specific enhancers). PHOX2B, GATA3 and HAND2 core regulatory circuit transcription factor ChIP-seq in a *MYCN*-expressing neuroblastoma cell line (green, CLB-GA).

### Local enhancer co-amplification explains asymmetric MYCN amplicon distribution

*MYCN* is expressed at the highest levels in neuroblastomas with *MYCN* amplifications (Supplementary Fig. 1d). It is unclear, however, to what extent enhancers are required for sustained MYCN expression on *MYCN*-containing amplicons. To address this, we mapped amplified genomic regions in a meta-dataset of copy-number variation in 240 *MYCN*-amplified neuroblastomas^21^. This revealed an asymmetric pattern of *MYCN* amplification (Fig. 2a, Supplementary Fig. 2). Intriguingly, a 290kb region downstream of *MYCN* was co-amplified in more than 90% of neuroblastomas, suggesting that *MYCN* amplicon boundaries were not randomly distributed, which is in line with recent reports in a smaller tumor cohort^19^ Notably, the consensus amplicon boundaries did not overlap with common fragile sites (Supplementary Fig. 2g), challenging a previous association found in ten neuroblastoma cell lines^8^. Regions of increased chromosomal instability alone are therefore unlikely to explain amplicon boundaries. Intriguingly, several *MYCN*-specific enhancers were found to be commonly co-amplified (Fig. 2b). The distal *MYCN*-specific CRC-driven enhancer, e4, was part of the consensus amplicon region in 90% of cases. Randomizing amplicon boundaries around *MYCN* showed that e4 coamplification was significantly enriched on *MYCN* amplicons (empirical *P*=0.0003). Coamplification frequency quickly dropped downstream of e4, suggesting that *MYCN*-specific, CRC-driven enhancers are a determinant of *MYCN* amplicon structure and may be required for *MYCN* expression, even in the context of high-level amplification.

**Fig. 2.**
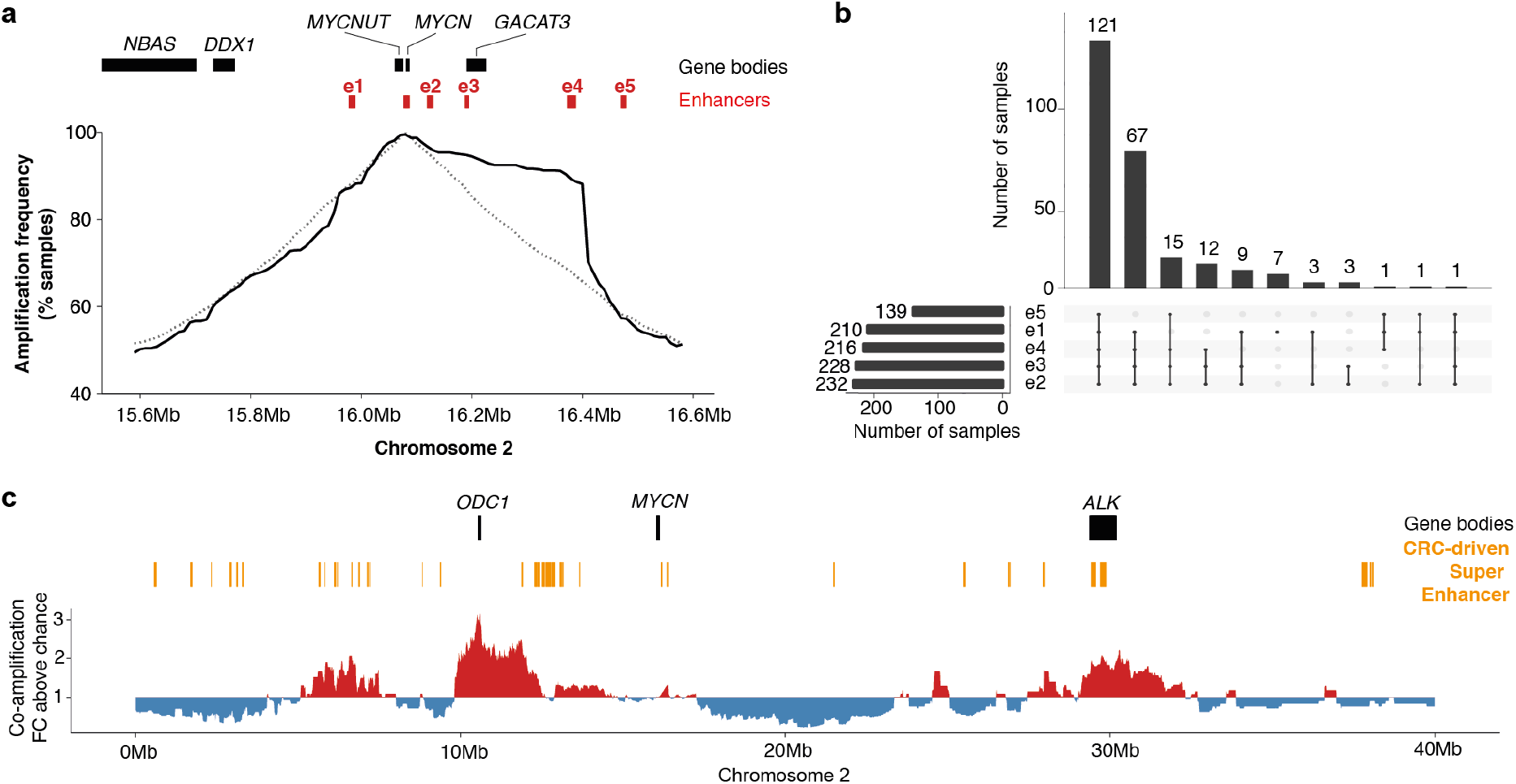
*MYCN*-specific enhancer e4 is significantly co-amplified with *MYCN* and retains functional enhancer characteristics after amplification. **a,** Co-amplification frequency of the immediate *MYCN* neighborhood measured using copy number profiles from 240 *MYCN*-amplified neuroblastomas (solid line) compared to the expected co-amplification frequencies for randomized *MYCN*-containing amplicons (dashed line). **b,** Upset plot showing the coamplification patterns of all five *MYCN*-specific local enhancers identified in neuroblastoma. **c,** Enrichment for co-amplification with *MYCN* of genomic regions on 2p (red, co-amplification more frequent than expected by chance; blue, co-amplification less frequent than expected by chance).

### Distal CRC-driven super enhancers are significantly co-amplified with MYCN in neuroblastoma

We and others have previously described chimeric *MYCN* amplicons^10^ containing distal chromosomal fragments. We therefore systematically inspected *MYCN*-distal regions on chromosome 2 for signs of co-amplification. Distinct regions were statistically enriched for coamplification with *MYCN* (Fig. 2c). In line with previous reports^22^, significant co-amplification of 19 protein-coding genes, including known neuroblastoma drivers such as *ODC1, GREB1* and *ALK* occurred in *MYCN*-amplified neuroblastoma. Intriguingly, co-amplification of distal CRC-driven super enhancers (SE) occurred in 23.3% of samples. Seven specific CRC-driven SEs were significantly co-amplified more often than expected by chance. Most of these SEs were found in gene-rich regions, precluding to determine whether genes or regulatory elements were driving co-amplification. One significantly co-amplified CRC-driven SE, however, was found in a gene-poor region in 2p25.2, where most co-amplified segments did not overlap protein-coding genes (Fig. 2c). This raised the question whether hijacking of such distal regulatory elements may explain co-amplification with *MYCN*.

### Enhancers remain functional on MYCN amplicons

Based on our amplicon boundary analysis, two classes of *MYCN* amplicons could be distinguished in neuroblastoma, i. amplicons containing local MYCN-specific enhancers, including e4, (here referred to as class I amplicons; Fig. 3a) and ii. amplicons lacking local *MYCN*-specific enhancers, and at least lacking e4 (referred to as class II amplicons; Fig. 3b). To determine whether co-amplified enhancers were active, we acquired genomic (long- and short-read whole genome sequencing) and epigenomic (ATAC-seq and H3K4me1 and H3K27ac ChIP-seq) data for two neuroblastoma cell lines with class I amplicons (Kelly and NGP) and two neuroblastoma cell lines with class II amplicons (IMR-5/75 and CHP-212). Notably, H3K27ac signal-to-noise ratio was lower on *MYCN* amplicons than in non-amplified regions. While the fraction of reads in peaks on the amplicon did not clearly differ between the amplicon and randomly drawn genomic regions, we observed more peaks than for nonamplified regions (Supplementary Fig. 3). These peaks were characterized by a lower relative signal compared to the amplicon background signal, indicating a larger variety of active regulatory regions on different *MYCN* amplicons. Using Nanopore long read-based *de novo* assembly, we reconstructed the *MYCN* neighborhood, confirming that *MYCN* and e4 were not only co-amplified in class I amplicons, but also lacked large rearrangements, which could preclude enhancer-promoter interaction (Supplementary Fig. 4-5). Enhancer e4 was characterized by increased chromatin accessibility and active enhancer histone marks as determined by ATAC-seq, H3K4me1 and H3K27ac ChIP-seq (Fig. 3c). Importantly, 4C chromatin conformation capture analysis showed that e4 spatially interacted with the *MYCN* promotor on the amplicon (Fig. 3c). Thus, e4 presents as a functional enhancer and appears to contribute to *MYCN* expression even in the context of class I *MYCN* amplification.

**Fig. 3.**
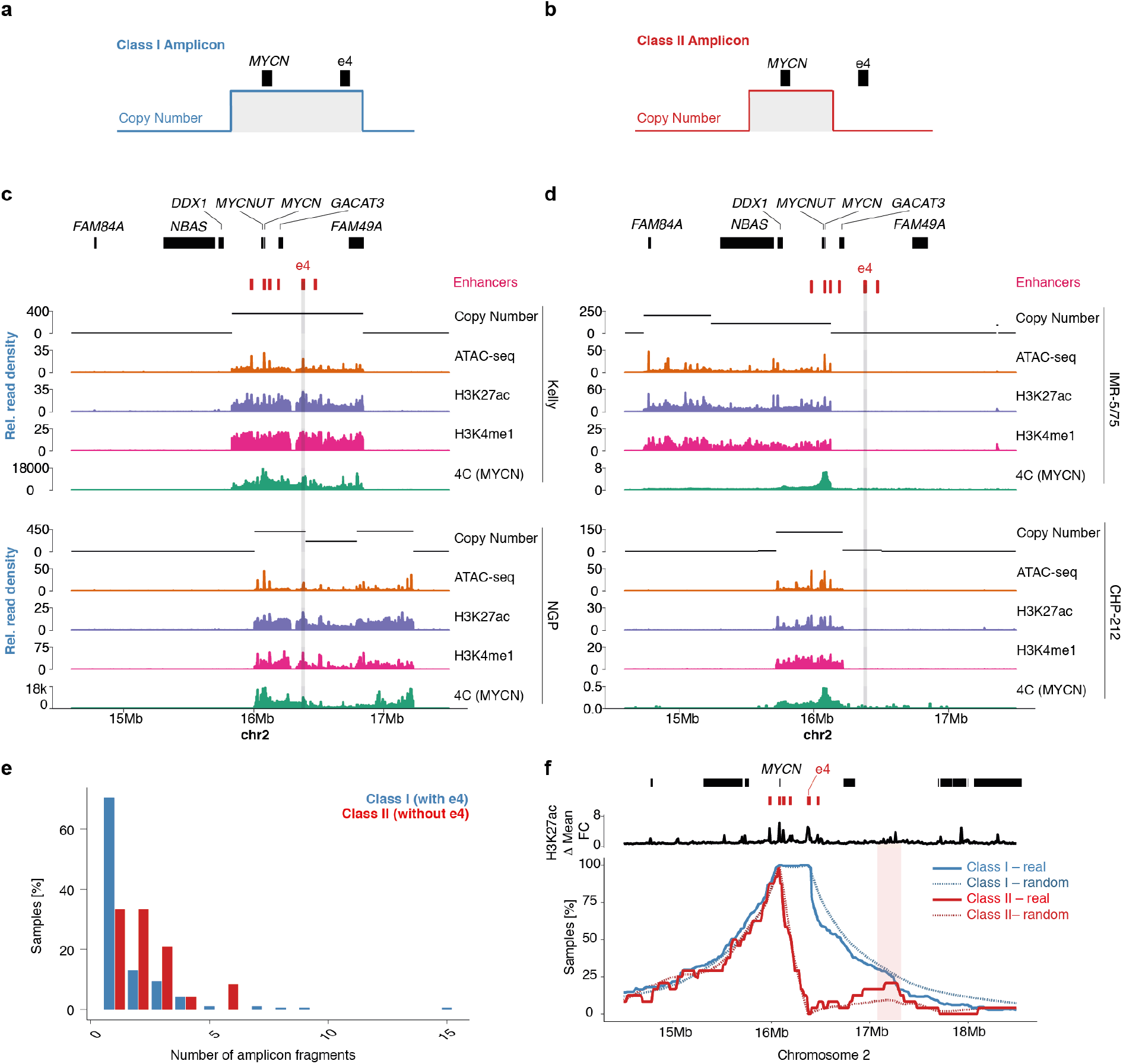
Two classes of *MYCN* amplicons can be identified in neuroblastoma. Schematic representation of class I (**a**) and class II (**b**) *MYCN* amplicons. **c,** Copy number profile, ATAC-seq, H3K27ac ChIP-seq, H3K4me1 ChIP-seq and 4C (*MYCN* promotor as the viewpoint) for two neuroblastoma cell lines with class I amplicons, co-amplifying the e4. **d,** Copy number profile, ATAC-seq, H3K27ac ChIP-seq, H3K4me1 ChIP-seq and 4C (*MYCN* promotor as the viewpoint) for two neuroblastoma cell lines class II amplicons, not co-amplifying e4. **e,** Number of non-contiguous amplified fragments in class I vs. class II *MYCN* amplicons. **d** Amplicon boundary frequency relative to gene and enhancer positions in class I (blue) vs. class II (red) amplicons compared to random amplicon boundary frequencies (dotted lines).

### Super enhancer hijacking compensates for the loss of local enhancers on chimeric intra- and extrachromosomal circular MYCN amplicons

In contrast to class I amplicons, class II amplicons did not include local enhancers, raising the possibility of alternative routes of *MYCN* regulation. The lack of a strong local regulatory element on class II amplicons and our observation of frequent co-amplification of distal SE (Fig. 2c), led us to hypothesize that ectopic enhancers might be recruited to enable *MYCN* expression in class II amplicons. In line with our hypothesis, primary neuroblastomas with class II amplicons were more likely to harbor complex amplifications containing more than one fragment (66.7% vs. 35.7%, Fisher’s Exact Test *P*=0.003; Fig. 3e). All class II amplicons coamplified at least one CRC-driven super enhancer element distal of *MYCN*. Some enhancers were recurrently found on class II amplicons, including an enhancer 1.2Mb downstream of *MYCN* that was co-amplified in 20.8% (5/24) of *MYCN*-amplified neuroblastomas, 2.1-fold higher than expected for randomized amplicons that include *MYCN* but not e4 (Fig. 3f). Thus, class II *MYCN* amplicons are of high chimeric structural complexity allowing for the replacement of local enhancers through hijacking of distal CRC-driven enhancers.

To determine the structure and epigenetic regulation of class II amplicons in detail, we inspected long-read based *de novo* assemblies and short read-based reconstructions of IMR-5/75 and CHP-212 *MYCN* amplicons. IMR-5/75 was characterized by a linear HSR class II *MYCN* amplicon, not including e3-e5 (Fig. 3b). Inspection of the IMR-5/75 *MYCN* amplicon structure revealed that the amplicon consisted of six distant genomic regions, which were joined together to form a large and complex chimeric amplicon (Fig. 4a-d). In line with enhancer hijacking, an intronic segment of *ALK* containing a large super enhancer, marked by H3K27ac modification and chromatin accessibility, was juxtaposed with *MYCN* on the chimeric amplicon. Similar to e4, this enhancer was bound by adrenergic CRC factors in non-amplified cells (Supplementary Fig. 6a). Notably, a CTCF-bound putative insulator was added to the amplicon by yet another distal fragment (Fig. 4a-c, Supplementary Fig. 6a). In CHP-212, *MYCN* is amplified on extrachromosomal circular DNA, as confirmed by fluorescence in situ hybridization (Supplementary Fig. 7). Both *de novo* assembly and short-read based reconstruction of the amplicon confirmed the circular *MYCN* amplicon structure independently (Fig. 4f-h). Similar to IMR-5/75, distal fragments containing CRC-driven SEs and putative CTCF-bound insulators were joined to the *MYCN* neighborhood (Fig. 4e-g, Supplementary Fig. 6b).

**Figure 4.**
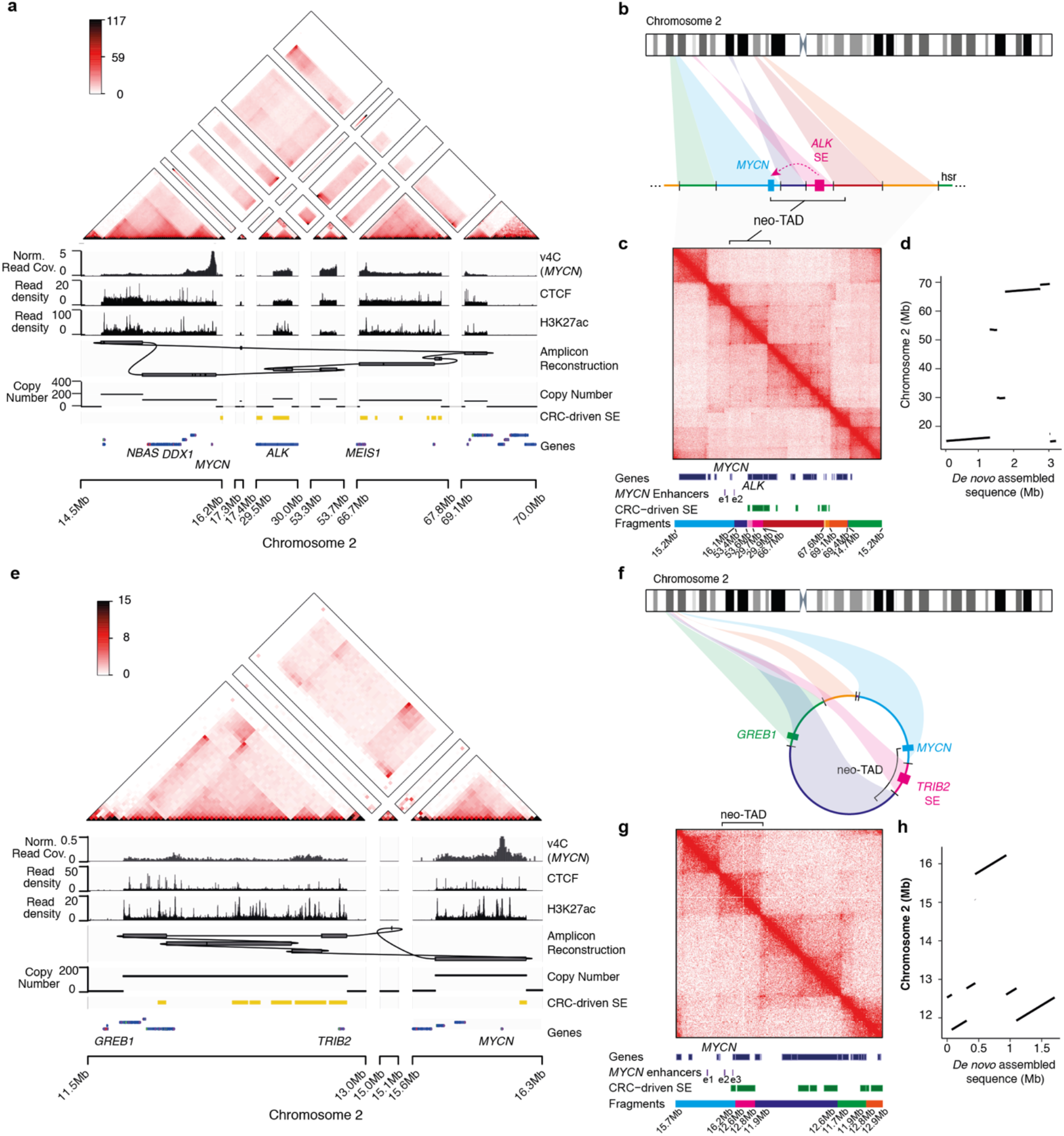
Reconstruction and epigenetic markup of class II intra- and extrachromosomal circular *MYCN* amplicons in neuroblastoma cells. **a** Short-read based reconstruction and epigenomic characterization of the *MYCN* amplicon in IMR-5/75 cells. Top to bottom: Hi-C map (color indicating Knight-Ruiz normalized read counts in 25kb bins), virtual 4C (MYCN viewpoint, v4C), CTCF ChIP-seq, H3K27Ac ChIP-seq, Amplicon reconstruction, copy number profile, super enhancer locations (yellow), gene positions (blue) (scale). **b** Schematic representation of the class II amplicon described in (**a**), showing ectopic enhancers and insulator reshuffling leading to locally disrupted regulatory neighborhoods on the HSR. **c** Alignment of Hi-C reads to the reconstructed *MYCN* amplicon in IMR-5/75 and positions of genes, local *MYCN* enhancers and CRC-driven super enhancers on the amplicon. **d** Mapping of the long read sequencing-based *de novo* assembly of the *MYCN* amplicon in IMR5/75 on chromosome 2. **e** Short-read based reconstruction and epigenomic characterization of the *MYCN* amplicon in CHP-212 cells. Top to bottom: Hi-C map (color indicating Knight-Ruiz normalized read counts in 25kb bins), virtual 4C (MYCN viewpoint, v4C), CTCF ChIP-seq, H3K27Ac ChIP-seq, Amplicon reconstruction, copy number profile, super enhancer locations (yellow), gene positions (blue). **f**, Schematic representation of the class II amplicon described in (**e**), showing ectopic enhancers and insulator reshuffling leading to locally disrupted regulatory neighborhoods on extrachromosomal circular DNA. **g** Alignment of Hi-C reads to the reconstructed *MYCN* amplicon in CHP-212 and positions of genes, local *MYCN* enhancers and CRC-driven super enhancers on the amplicon. **h** Mapping of the long read sequencing-based *de novo* assembly of the *MYCN* amplicon in CHP-212 on chromosome 2.

To analyze the interaction profile in circular and linear amplicons we performed Hi-C and mapped the reads to the reconstructed amplicon (Fig. 4c, g). This analysis supported the genomic sequencing-based reconstruction of the amplicon, recapitulating the order and orientation of the joined fragments and confirmed that the ectopic enhancers spatially interacted with *MYCN*. Notably, high-frequency interactions in the corners of the maps opposite to the main diagonal, confirmed the circularity of CHP-212 amplicon and the presence of tandem amplification in IMR-5/75. In IMR-5/75 and CHP-212, we observed insulated TADs, boundaries and loops as in the rest of the genome. Due to the rearrangements in CHP-212, the *MYCN* gene became part of a neo-TAD consisting of a sub-TAD that originated from the wild type genome as an intact unit, and a second sub-TAD that resulted from the fusion and coamplification of the first region with another region from a distal part of chromosome 2 (chr2:12.6-12.8Mb), containing multiple CRC-driven SEs (Fig. 4g, Supplementary Fig. 6b). Since the fused segments are now part of one TAD and not separated by a boundary, *MYCN* interaction with the SEs in this region becomes possible. A similar situation was observed for the linear amplicon. In IMR-5/75, Hi-C showed frequent contacts between *MYCN* and SEs from the genomic regions juxtaposed to *MYCN*, containing intronic parts of *ALK* (Fig. 4c, Supplementary Fig. 6a). The map also reflected the high complexity and genomic heterogeneity of the IMR-5/75 amplicon. Nevertheless, the TAD structure, boundaries and loops were clearly visible on the reconstructed Hi-C map. Thus, hijacking of ectopic enhancers and insulators can compensate for the loss of endogenous regulatory elements on intra- and extrachromosomal circular *MYCN* amplicons via the formation of neo-TADs, which may explain the higher structural complexity of *MYCN* amplicons lacking endogenous enhancers.

### Nanopore long-read DNA sequencing can be used for parallel assessment of MYCN amplicon structure and epigenetic regulation

In addition to allowing the alignment-free *de novo* assembly of the *MYCN* amplicon in several samples (Fig. 4b-d, f-h, Supplementary Fig. 4-5), Nanopore sequencing also allows for the direct measurement of DNA methylation without the need for bisulfite conversion (Fig. 5a)^23^. While DNA methylation at regulatory elements is often associated with repression, a trough in DNA methylation may indicate a transcription factor binding event, a poised or active gene regulatory element, or a CTCF-occupied insulator element (Fig. 5b). In theory, Nanopore sequencing and assembly might allow for the simultaneous inference of both structure and regulatory landscape (Fig. 5b). Prior to evaluating the *MYCN* amplicons, the DNA methylation landscape of highly expressed and inactive genes demonstrated the expected distribution of decreased methylation at active promoters and increased methylation within active gene bodies (Fig. 5c). In order to assess the DNA methylation status of putative regulatory elements near *MYCN*, we first used the amplicon-enriched ATAC-seq peaks to classify relevant motif signatures (Fig. 5d). While *MYCN* was surrounded by the expected CRC-driven regulatory elements at the overlapping core enhancers as well as some CTCF sites, both their number and location varied, indicative of sample-specific sites of regulation. Indeed, DNA methylation decreased in accordance with sites specific to a given sample (Fig. 5e), opening up the possibility of using these data to infer regulatory elements in patient samples when no orthogonal epigenomic data are available.

**Figure 5.**
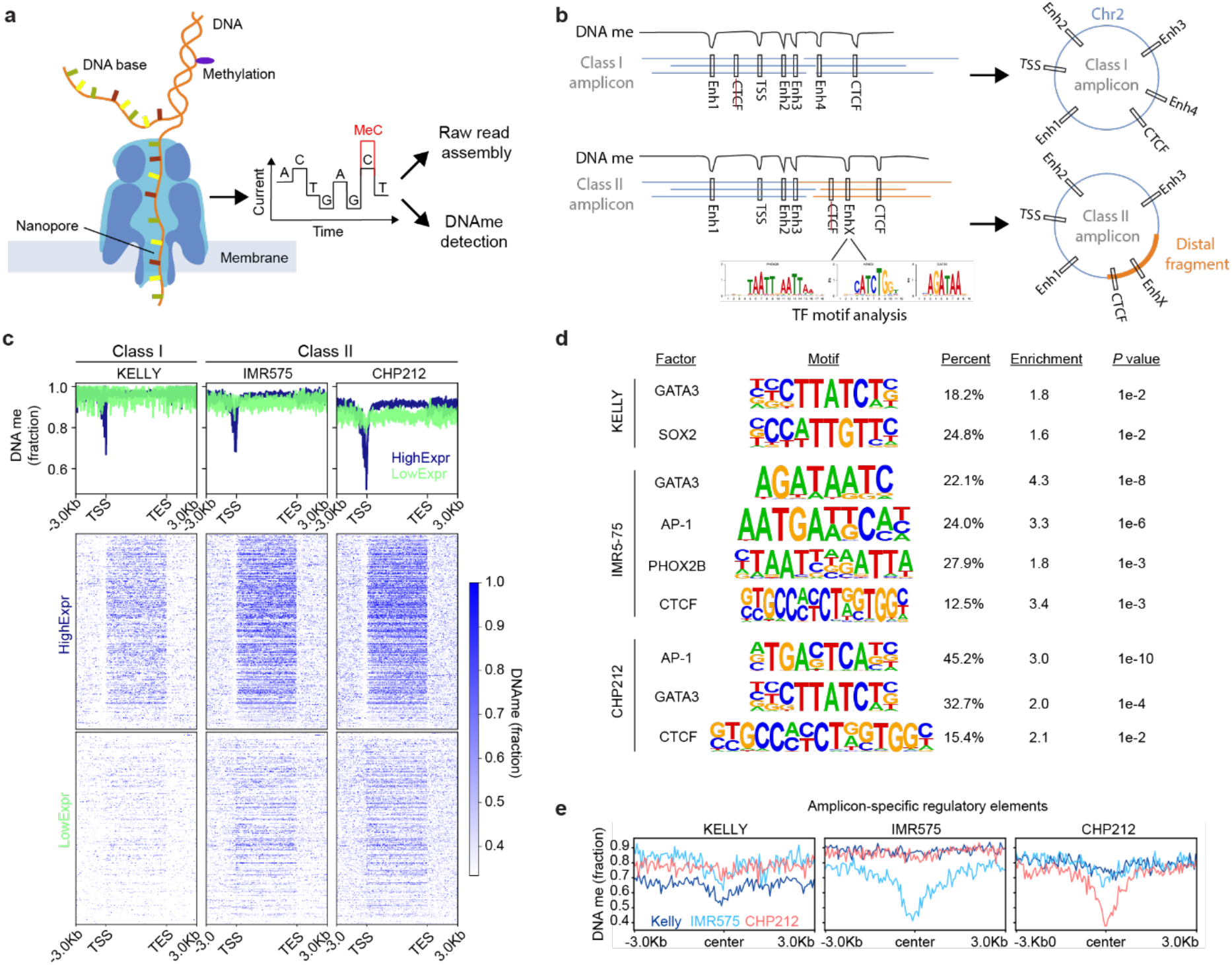
Nanopore long read sequencing allows for the simultaneous characterization of amplicon structure and DNA methylation. **a,** Schematic of experimental approach. **b,** Schematic representation of how Nanopore sequencing facilitates *de novo* amplicon assembly and can be used to simultaneously to detect regulatory elements through DNA methylation analysis. **c,** Composite DNA methylation signal detected using Nanopore sequencing over genes expressed at high (blue) vs. low (green) levels. **d,** Motif analysis based on accessibility in regulatory elements co-amplified on *MYCN* amplicons. **e,** Amplicon-specific methylation pattern detected in three neuroblastoma cell lines using Nanopore sequencing-based DNA methylation analysis.

### Class II MYCN amplicons clinically phenocopy class I amplicons

*MYCN*-amplified neuroblastoma is characterized by significant clinical heterogeneity, which cannot entirely be explained genetically. Whether the structure of the *MYCN* amplicon itself could account for some of this variation is currently unknown. In line with previous reports^22^, higher counts of amplified fragments were associated with a more malignant clinical phenotype (Fig. 6a). Co-amplification of *ODC1*, a gene located 5.5Mb upstream of *MYCN* and coamplified in 9% (21/240) of *MYCN*-amplified neuroblastomas (Fig. 2c), defined an ultra-high risk genetical subgroup of *MYCN*-amplified neuroblastoma (HR 2.3 (1.4-3.7), Log-rank test *P*=0.001; Fig. 6b). Similarly, *ALK* co-amplification, present in in 5% (12/240) of *MYCN*-amplified tumors, was also associated with adverse clinical outcome (HR 1.8 (0.94-3.4), Logrank test *P*=0.073; Fig. 6c). In contrast, differences in the *MYCN* amplicon enhancer structure, i.e. class II amplification, did not confer prognostic differences (HR 1.3 (0.78-2.1), Log rank test *P*=0.34; Fig. 6d). We therefore conclude that chimeric co-amplification of proto-oncogenes partly explain the malignant phenotype of neuroblastomas with complex *MYCN* amplicons, whereas enhancer hijacking in class II amplicons does not change clinical behavior, fully phenocopying class I *MYCN* amplicons.

**Figure 6.**
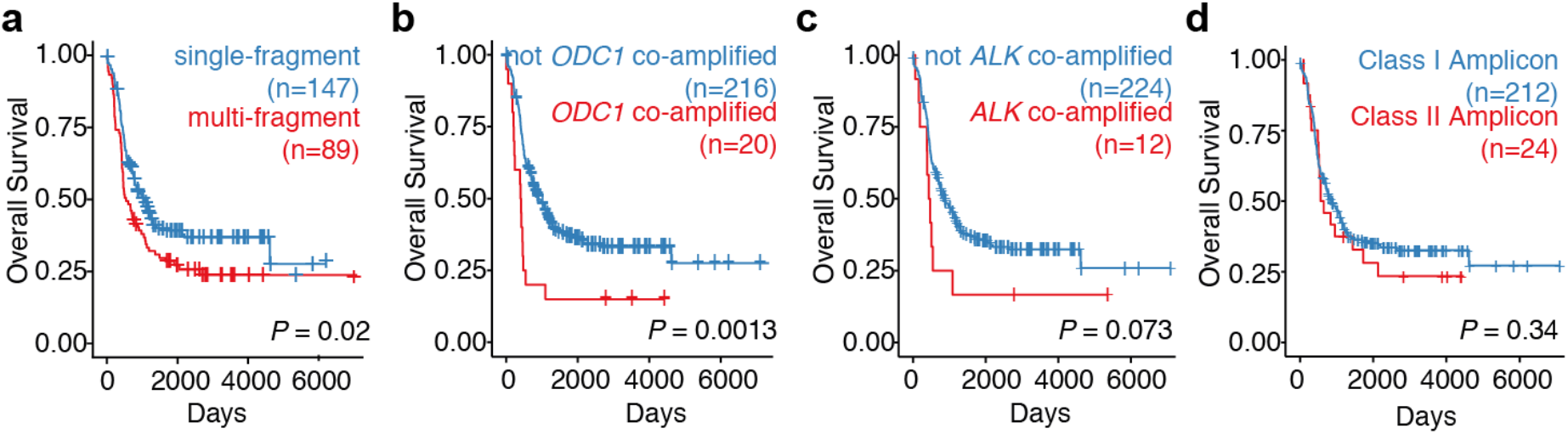
Class II amplicons clinically phenocopy class I amplicons. Kaplan Meier survival analysis of patients with *MYCN*-amplified neuroblastoma, comparing single-fragment vs. non-contiguously amplified *MYCN* amplicons (**a**), co-amplification of *ODC1* vs. no coamplification (**b**), co-amplification of *ALK* vs. no co-amplification, and class I amplicons vs. class II amplicons (**d**; *N*=236 *MYCN*-amplified neuroblastomas; *P*-value based on two-sided log rank test).

## Discussion

Here, we show that neuroblastoma-specific CRC-driven enhancers contribute to *MYCN* amplicon structure in neuroblastoma and retain the classic features of active enhancers after genomic amplification. While most *MYCN* amplicons contain local enhancers, ectopic enhancers are regularly incorporated into chimeric amplicons lacking local enhancers, leading to enhancer hijacking.

A large subset of neuroblastomas was recently found to be driven by a small set of transcription factors that form a self-sustaining core regulatory circuit, defined by their high expression and presence of super-enhancers^15–18^. In how far *MYCN* itself is directly regulated by CRC factors was previously unclear, particularly due to the challenging interpretation of epigenomic data on amplicons^16^. Our results provide empiric evidence that *MYCN* is driven by CRC factors even in the context of *MYCN* amplification. This is in line with and can mechanistically explain the previous observation that genetic depletion of CRC factors represses *MYCN* expression even in *MYCN*-amplified cells^16^. The finding that ectopic enhancers driven by the CRC are juxtaposed to *MYCN* on amplicons that lack local enhancers further strengthens the relevance of the CRC in *MYCN* regulation.

In line with our observation of local enhancer co-amplification, Morton et al. recently described that local enhancers are significantly co-amplified with other proto-oncogenes in other cancer entities^19^. They showed that experimentally interfering with local *EGFR* enhancers in *EGFR*-amplified glioblastoma impaired oncogene expression and cell viability in *EGFR*-amplified as well as non-amplified cases. In line with our findings, a region overlapping e4 was identified to be significantly co-amplified in *MYCN*-amplified neuroblastomas, corresponding to class I amplicons observed in our cohort. In contrast to Morton et al., who suggest that the inclusion of local enhancers is necessary for proto-oncogene expression on amplicons, we show that exceptions to this rule occur in a significant subset of *MYCN* amplified neuroblastomas. In such cases, amplicons are of highly complex chimeric structure enabling the reshuffling of ectopic enhancers and insulators to form neo-TADs that can compensate for disrupted local neighborhoods through enhancer hijacking.

More generally, we show that TADs also form in ecDNA, in line with recent findings by Wu et al.^24^. We extend this observation to homogeneously staining regions, which form extremely expanded stretches of chromatin in interphase nuclei and lose chromosomal territoriality^25^. Gene activation by enhancer adoption requires the fusion of distant DNA fragments and the formation of new chromatin domains, called neo-TADs^26^. This fusion requires a convergent directionality of CTCF sites in order to form a new boundary. Only in this case, aberrant gene activation is possible^27^.

Reconstruction of amplicons has previously relied on combining structural breakpoint coordinates to infer the underlying structure. This regularly resulted in ambiguous amplicon reconstructions, which had to be addressed by secondary data such as Chromium linked reads or optical mapping^4,6,24^. We demonstrate the feasibility of long-read *de novo* assembly for the reconstruction of amplified genomic neighborhoods. *De novo* assembly was able to reconstruct entire ecDNA molecules and confirm the tandem duplicating nature of homogeneously staining regions. Integrating *de novo* assembly with methylation data from Nanopore sequencing reads will likely benefit further studies of other proto-oncogene-containing amplicons by enabling the characterization of the interplay between structure and regulation in highly rearranged cancer genomes.

Functional studies have shown that both *ODC1* and *ALK* are highly relevant in neuroblastoma ^28,29^. Co-amplification with *MYCN* has been reported before^22^, but to our knowledge the clinical relevance of co-amplification had not been determined so far. Similar to our previous observations of *PTP4A2* co-amplification on chimeric ecDNA^10^, we demonstrate here that proto-oncogenes reside side-by-side on the same extrachromosomal circular DNAs, sometimes even sharing the same regulatory neighborhood. It is tempting to speculate that this structural coupling of genes could confer MYCN-independent but *MYCN*-amplicon-specific, collateral therapeutic vulnerabilities in *MYCN*-amplified tumors.

We conclude that the structure of genomic amplifications can be explained by selective pressure not only on oncogenic coding elements, but also on non-coding regulatory elements. CRC-driven enhancers are required for successful *MYCN* amplification and remain functional throughout this process. Even though the majority of amplicons contain endogenous enhancers, these can be replaced by ectopic CRC-driven elements that are juxtaposed to the oncogene through complex chimeric amplicon formation. We envision that our findings also extend to oncogene amplifications in other cancers and will help identify functionally relevant loci amongst the diverse array of complex aberrations that drive cancer.

## Materials and Methods

### Cell lines

Neuroblastoma cell lines (CHP-212, IMR-5/75, NGP, Kelly) were a gift from from Carol J. Thiele, obtained from the German Collection of Microorganisms and Cell Cultures or obtained from the American Type Culture Collection. Cell line identity was verified by STR genotyping (Genetica DNA Laboratories, Burlington, NC and IDEXX BioResearch, Westbrook, ME) and absence of *Mycoplasma sp*. contamination was determined with a Lonza MycoAlert system (Lonza Group Ltd., Basel, CH). All cell lines were cultured in RPMI-1640 medium (Thermo Fisher Scientific, Inc., Waltham, MA) with 1% Penicillin/Streptomycin, and 10% FCS.

### RNA-seq

Public RNA-seq data was downloaded from Gene Expression Omnibus (GSE90683)^15^. FASTQ files were quality controlled (FASTQC 0.11.8) and adapters were trimmed (BBMap 38.58). We mapped reads to GRCh37 (STAR 2.7.1 with default parameters), counted them per gene (Ensembl release 75, featureCounts from Subread package 1.6.4) and normalized for library size and composition (sizeFactors from DESeq2 1.22.2).

### ChIP-seq

As reported before^27^, cells were digested with Trypsin–EDTA 0.05% (Gibco) for 10 min at 37 °C. The cells were mixed with 10% FCS–PBS, and a single-cell suspension was obtained using a 40-μm cell strainer ^30^. After centrifugation, cells were resuspended in 10% FCS-PBS again and fixed in 1% paraformaldehyde (PFA) for 10 min at room temperature and reaction quenched with 2.5M glycine (Merck) on ice and centrifuged at 400g for 8min. Pelleted cells were then resuspended in lysis buffer (50 mM Tris, pH 7.5; 150 mM NaCl; 5 mM EDTA; 0.5% NP-40; 1.15% Triton X-100; protease inhibitors (Roche)), and nuclei were pelleted again by centrifugation at 750g for 5min. For sonication, nuclei were resuspended in sonication buffer (10 mM Tris–HCl, pH 8.0; 100 mM NaCl; 1 mM EDTA; 0.5 mM EGTA; 0,1% Na-deoxycholate; 0.5%*N*-lauroylsarcosine; protease inhibitors (Roche complete)). Chromatin was sheared using a Bioruptor until reaching a fragment size of 200–500 base pairs (bp). Lysates were clarified from sonicated nuclei, and protein–DNA complexes were immunoprecipitated overnight at 4 °C with the respective antibody. A total of 10–15 μg chromatin was used for each replicate of histone ChIP and 20-25μg of transcription factor ChIP. Anti-H3K27ac (Diagenode; c15410037; A1657D), anti-H3K4m1 (Abcam; ab8895; Lot A1657D), anti-RAD21 (Abcam; ab992; Lot GR221348-8) and anti-CTCF (Active Motif; 613111; Lot 34614003) antibodies were used. Sequencing libraries were prepared using standard Nextera adapters (Illumina) according to the supplier’s recommendations. 25 million reads per sample were sequenced on HiSeq 2500 sequencer (Illumina) in 50bp single read mode.

Additional public ChIP-seq FASTQ files were downloaded from Gene Expression Omnibus (GSE90683, GSE24447 and GSE28874)^15,31^. FASTQ files were quality controlled (FASTQC 0.11.8) and adapters were trimmed (BBMap 38.58). Reads were then aligned to hg19 (BWA-MEM 0.7.15 with default parameters) and duplicate reads removed (Picard 2.20.4). We generated BigWig tracks by extending reads to 200bp for single-end libraries and extending to fragment size for paired-end libraries, filtering by ENCODE DAC blacklist and normalizing to counts per million in 10bp bins (Deeptools 3.3.0). Peaks were called using MACS2 (2.1.2) with default parameters. Super enhancers were called for H3K27ac data using LILY (https://github.com/BoevaLab/LILY) with default parameters. ChIP-seq data was quality controlled using RSC and NSC (Phantompeakqualtools 1.2.1).

### ATAC-seq

ATAC-seq samples were processed as reported in Buenrostro et al^32^. 5×10^5^ cells were used per sample. For sequencing, libraries were generated using Illumina/Nextera adapters and size selected (100–1000bp) with AMPure Beads (Beckman Coulter). Approximately 100 million 75bp paired-end reads were acquired per sample on the HiSeq 2500 system (Illumina). Additional public ATAC-seq FASTQ files were downloaded from Gene Expression Omnibus (GSE80154)^33^. Adapter trimming, alignment and duplicate removal as for ChIP-seq. We generated BigWig tracks by extending paired-end reads to fragment size, filtering by the ENCODE DAC blacklist and normalizing to counts per million in 10bp bins (Deeptools 3.3.0). Peaks were called using MACS2 (2.1.2) with default parameters.

### Hi-C

3C libraries for Hi-C and 4C were prepared from confluent neuroblastoma cells according to the cell culture section above. Hi-C experiments were performed as duplicates. Cells were washed twice with PBS and digested with Trypsin–EDTA 0.05% (Gibco) for 10 min at 37 °C. To obtain a single cell suspension, cells were pipetted through a 40-μm cell strainer ^30^.

After centrifugation at 300g for 5min, cell pellets were resuspended with 10% FCS and fixed by adding an equal volume of 4% formaldehyde (Sigma-Aldrich) and mixed for 10 min at room temperature while shaking. Fixation was quenched using 1.425 M glycine (Merck) on ice and immediately centrifuged at 400g for 8 min. Pelleted cells were then resuspended in lysis buffer (50 mM Tris, pH 7.5; 150 mM NaCl; 5 mM EDTA; 0.5% NP-40; 1.15% Triton X-100; protease inhibitors (Roche)), and nuclei were pelleted again by centrifugation at 750g for 5min.

The pellet was washed with 1x DpnII buffer, resuspended in 50μl 0.5% SDS and incubated for 10min at 62°C. After that 145μl water and 25μl 10% Triton (Sigma) was added to quench the SDS. After a 37°C incubation, 25μl DpnII buffer and 100U DpnII was added. The digestion reaction was incubated for 2h at 37°C, after 1h another 10U were added. After the digestion, DpnII was inactivated at 65°C for 20min.

The digested sticky ends were filled up with 10mM dNTPs (without dATP) and 0.4mM biotin-14-dATP (Life Technologies) and 40U DNA Pol I, Large Klenow (New England BioLabs, Inc. (NEB), Ipswich, MA) at 37°C for 90min. Biotinylated blunt ends were then ligated using a ligation reaction (663μl water, 120μl 10X NEB T4 DNA ligase buffer (NEB), 100μl 10% Triton X-100 (Sigma), 12μl 10mg/ml BSA and 2400U of T4 DNA liagse (NEB)) overnight at 16°C with slow rotation.

The 3C library was then sheared using a Covaris sonicator (duty cycle: 10%; intensity: 5; cycles per burst: 200; time: 6 cycles of 60 s each; set mode: frequency sweeping; temperature: 4–7 °C). After sonication, religated DNA was pulled down using 150μl of 10mg/ml Dynabeads Streptavidin T1 beads (Thermo Fisher) according to the supplier’s recommendation. Sheared and pulled down DNA was treated using a 100μl end-repair reaction (25mM dNTPs, 50U NEB PNK T4 Enzyme, 12U NEB T4 DNA polymerase, 5U NEB DNA pol I, Large (Klenow) Fragment, 10X NEB T4 DNA ligase buffer with 10mM ATP) and incubated for 30min at 37°C. Universal sequencing adaptor were added using the NEBnext Ultra DNA Library Kit (NEB) according to the supplier’s recommendation. Samples were sequenced with Ilumina Hi-Seq technology according to standard protocols and 75bp PE mode. 200 million reads were generated for IMR-5/75, 5 million reads per sample were generated for all other cell lines.

FASTQ files were processed using the Juicer pipeline v1.5.6, CPU version^34^, which was set up with BWA v0.7.17^35^ to map short reads to reference genome hg19, from which haplotype sequences were removed and to which the sequence of Epstein-Stein-Barr Virus (NC_007605.1) was added. Replicates were processed individually. Mapped and filtered reads were merged afterwards. A threshold of MAPQ≥30 was applied for the generation of Hi-C maps with Juicer tools v1.7.5^34^. Knight-Ruiz normalization of Hi-C signal was used for Hi-C maps. Virtual 4C signal for the *MYCN* locus was generated by the mean Knight-Ruiz-normalized Hi-C signal across three 5kb bins (chr2:16,075,000-16,085,000).

### 4C-seq

4C-seq libraries were generated as described before^26^, using a starting material of 5×10^6^ – 1×10^7^cells. The fixation and lysis were performed as described in the Hi-C section. For the *MYCN*promotor viewpoint, 1.6 μg DNA was amplified by PCR (Primer 1 5’-GCAGAATCGCCTCCG-3’, Primer 2 5’-CCTGGCTCTGCTTCCTAG-3’). For the viewpoint, 4bp cutters were used. DpnII (NEB) was used as first cutter and Csp6I (NEB) as second cutter. All samples were sequenced with the HiSeq 2500 (Illumina) technology according to standard protocols and with 8 million reads per sample.

Reads were pre-processed, filtered for artefacts and mapped to the reference genome GRCh37 using BWA-MEM as described earlier^26^. After removing the viewpoint fragment as well as 1.5 kb up- and downstream of the viewpoint the raw read counts were normalized per million mapped reads (RPM) and a window of 10 fragments was chosen to smooth the profile.

### Whole-genome sequencing

Cells were harvested and DNA was extracted using the NucleoSpin Tissue kit (Macherey-Nagel GmbH & Co. KG, Düren, Germany). Libraries for whole genome sequencing were prepared with the NEBNext Ultra II FS DNA Library Prep Kit for Illumina (New England BioLabs, Inc., Ipswich, MA). Libraries were sequenced on a NovaSeq S1 flow cell (Illumina, Inc., San Diego, CA) with 2×150bp paired-end reads. Quality control, adapter trimming, alignment, duplicate removal as for ChIP-seq data. Copy number variation was called (Control-FREEC^36^ 11.4 with default parameters). Structural variants were called using SvABA^37^ (1.1.1) in germline mode and discarding regions in a blacklist provided by SvABA (https://data.broadinstitute.org/snowman/svaba_exclusions.bed).

### Nanopore Sequencing

Cells were harvested and high molecular weight DNA was extracted using the MagAttract HMW DNA Kit (Qiagen N.V., Venlo, Netherlands). Size selection was performed to remove fragments <10 kilobases (kb) using the Circulomics SRE kit (Circulomics Inc., Baltimore, MD). DNA content was measured with a Qubit 3.0 Fluorometer (Thermo Fisher) and sample quality control was performed using a 4200 TapeStation System (Agilent Technologies, Inc., Santa Clara, CA). Libraries were prepared using the Ligation Sequencing Kit (SQK-LSK109, Oxford Nanopore Technologies Ltd., Oxford, UK) and sequenced on a R9.4.1 MinION flowcell (FLO-MIN106, Oxford Nanopore Technologies Ltd., Oxford, UK). Quality control was performed using NanoPlot 1.0.0. For the NGP cell line, DNA was extracted with the NucleoSpin Tissue kit (Macherey-Nagel GmbH & Co. KG, Düren, Germany) and libraries prepared using the ONT Rapid Kit (SQK-RBK004, Oxford Nanopore Technologies Ltd., Oxford, UK). Guppy 2.3.7 (Oxford Nanopore Technologies Ltd., Oxford, UK) was used for basecalling with default parameters. For de novo assembly, Flye 2.4.2^38^ was run in metagenomics assembly mode on the unfiltered FASTQ files with an estimated genome size of 1Gb. Contigs were mapped back to hg19 using minimap2 2.16 with parameter -ax asm5. Assembly results were visualized with Bandage 0.8.1 (https://rrwick.github.io/Bandage) and Ribbon (no version available, https://github.com/MariaNattestad/Ribbon). CpG methylation was called from the unfiltered raw FAST5 files using Megalodon 0.1.0 (Oxford Nanopore Technologies Ltd., Oxford, UK).

### Fluorescence in situ hybridization

Cells were grown to 200,000 per well in six-well plates and metaphase-arrested using Colcemid (20μl/2ml; Roche #10295892001) for 30min-3h, trypsinized, centrifuged (1000rpm/10min) washed and pelleted. 5ml 0.4% KCl (4°C; Roth #6781.1) was added to the pellet and incubated for 10min. 1ml KCl and 1ml MeOH/acetic acid 3:1 (Roth #4627.2, #KK62.1) was added dropwise. 2/5/5ml of MeOH/acetic acid were added in between centrifugation steps (1000 rpm/10min) respectively. Suspension was dropped on a slide from a height of 40cm. Slides were washed with PBS (Gibco, #70011036) and digested for 10min in 0,04% pepsin solution in 0,001N HCl. Slides were washed in 0.5x SSC, dehydrated with 70%/80%/100% EtOH (3min each) and air-dried. 10μl of the probe (Vysis LSI N-MYC; #07J72-001; Lot #472123; Abbott Laboratories, Abbott Park, IL) were added and coverslips fixed on the slide. Slides were incubated at 75°C for 10min and at 37°C over night. The coverslip was removed and the slide washed in 0.4xSSC/0.3% IGEPAL (CA-630, #18896, Sigma-Aldrich Inc.) for 3min at 60°C and 2xSSC/0.1% IGEPAL for 3min at RT. 5μl DAPI (Vectashield, #H-1200, Vector) was added. A coverslip was added and fixed with nail polish.

### Enhancer calling

*MYCN*-expressing cell lines were defined as cell lines with sizeFactor normalized expression of 100 or above based. We identified enhancer candidate regions in a ±500kb window around *MYCN*. We focused on regions with a H3K27ac peak in the majority of *MYCN*-expressing, non-*MYCN*-amplified cell lines, i.e. three or more. If the gap between two such regions was less than 2kb, they were joined. These regions were then ranked by the maximum difference in H3K27ac signal fold change between non-amplified, *MYCN*-expressing and non-expressing cell lines. We chose the five highest-ranking regions as candidate regulatory elements. Enhancer regions were screened for transcription factor binding sequences from the JASPAR2018 (http://jaspar2018.genereg.net/) and JASPAR2020 (http://jaspar2020.genereg.net/) database using the TFBSTools (1.20.0) function matchPWM with min.score=‘85%’. CRC-driven super enhancers were defined as all regions with a LILY-defined super enhancer in *MYCN*-expressing, non-*MYCN*-amplified cell lines that overlapped with a GATA3, HAND2 or PHOX2B peak in CLB-GA.

### Analysis of copy number data

Public data was downloaded. Samples that were described as *MYCN*-amplified in the metadata but did not show *MYCN* amplification in the copy number profile were excluded. In order to generate an aggregate copy number profile, the genome was binned in 10kb bins and number of samples with overlapping amplifications was counted per bin. Randomized copy number profiles were generated by randomly sampling one of the original copy number profiles on chromosome 2 and randomly shifting it such that *MYCN* is still fully included within an amplified segment. For class I-specific shuffling, e4 had to be included as well; for class II-specific shuffling, e4 was never included on the randomly shifted amplicon. Empirical *P*-values for significant co-amplification were derived by creating 10,000 randomized datasets with each amplicon randomly shifted and comparing the observed co-amplification frequency to the distribution of co-amplification frequencies in the randomized data. Empirical *P*-values were always one-sided and adjusted for multiple comparisons using the Benjamini-Hochberg procedure.

### Amplicon reconstruction

All unfiltered SvABA structural variant calls were filtered to exclude regions from the ENCODE blacklist^39^ and small rearrangements of 1kb or less. As we were only aiming at the rearrangements common to all amplicons, we only considered breakpoints with more than 50 variant-support reads (‘allele depth’). gGnome^40^ was used to represent these data as a genome graph with nodes being breakpoint-free genomic intervals and edges being rearrangements (‘alternate edge’) or connections in the reference genomes (‘reference edge’). We considered only nodes with high copy number, i.e. with a mean whole-genome sequencing coverage of at least 10-fold the median coverage of chromosome 2. Then, reference edges were removed if its corresponding alternate edge was among the 25% highest allele-depth edges. The resulting graph was then searched for the circular, *MYCN*-containing walk that included the highest number of nodes without using any node twice. We used gTrack (https://github.com/mskilab/gTrack) for visualization. For custom Hi-C maps of reconstructed amplicon sequences of CHP-212 and IMR-5-75, respectively, the corresponding regions from chromosome 2 were copied, ordered, oriented and compiled according to the results from the amplicon reconstruction and added to the reference genome. Additionally, these copied regions were masked with ‘N’ at the original locations on chromosome 2 to allow a proper mapping of reads to the amplicon sequence. The contribution of Hi-C di-tags from these regions on chromosome 2 to the amplicon Hi-C map is expected be minor, because the copy number of amplicons is much higher than the number of wild type alleles. Juicebox v1.11.08 was used to visualize Hi-C maps with a bin size of 5 kb and Knight-Ruiz normalization^41–43^.

## Data availability

Copy number data for high-risk neuroblastoma were downloaded from https://github.com/padpuydt/copynumber_HR_NB/. Sequencing data supporting the findings of this manuscript is available at the Gene Expression Omnibus under accessions GSE90683, GSE80152, GSE24447 and GSE28874. Sequencing data for primary neuroblastoma samples is available at the European Genome-Phenome archive under accessions EGAS00001001308 and EGAS00001004022. Corresponding BigWig und narrowPeak files can be downloaded from https://data.cyverse.org/dav-anon/iplant/home/konstantin/helmsaueretal/. An accompanying UCSC genome browser track hub is provided for ChIP-seq and ATAC-seq data visualization (https://de.cyverse.org/dl/d/27AA17DA-F24C-4BF4-904C-62B539A47DCC/hub.txt). All other data is available from the corresponding authors upon reasonable request.

## Code availability

Code is available at https://github.com/henssenlab/MYCNAmplicon.

## Acknowledgments

We thank the patients and their parents for granting access to the tumor specimen and clinical information that were analyzed in this study. We are grateful to Yingqian Zhan, Natalia Munoz Perez, Jennifer von Stebut, Victor Bardinet and Celine Chen for critical discussions. R.P.K is supported by the Berlin Institute of Health visiting professorship program. A.G.H. is supported by the Deutsche Forschungsgemeinschaft (DFG, German Research Foundation) – 398299703, the Wilhelm Sander Stiftung a participant in the BIH-Charité Clinical Scientist Program funded by the Charité – Universitätsmedizin Berlin and the Berlin Institute of Health. A.G.H. and K.H. are supported by Berliner Krebsgesellschaft e.V. K.H. is supported by Boehringer Ingelheim Funds. This work was also supported by the TransTumVar project – PN013600. We thank B. Hero, H. Düren, N. Hemstedt of the neuroblastoma biobank and neuroblastoma trial registry (University Children’s Hospital Cologne) of the German Society of Pediatric Oncology and Hematology (GPOH) for providing samples and clinical data.

## Author Contributions

All authors contributed to the study design and collection and interpretation of the data. M.V. and S.A. acquired ChIP-seq, ATAC-seq, 4C and Hi-C data. R.C., L.P.K. and P.E. acquired nanopore sequencing data. K.K. acquired Illumina whole genome sequencing data. C.Rö. and C.Ro. performed FISH experiments. R.S., V.H. and K.H. analyzed 4C and Hi-C data. K.H. analyzed ChIP-seq, ATAC-seq and RNA-seq data. E.F., M.P., J.T. and K.H. analyzed Illumina whole-genome sequencing data. K.H. and R.P.K. analyzed nanopore sequencing data. M.F., F.H., A.K.-S. and J.H.S. collected and prepared patient samples. M.V., S.A., R.C., Y.B., H.D.G. and K.Ha. performed experiments and analyzed data. K.Ha., M.R., D.T. and J.H.S. contributed to study design. K.H., M.V., S.A., S.M., A.G.H. and R.P.K. led the study design, performed data analysis and wrote the manuscript, to which all authors contributed.

## Materials & Correspondence

Request for materials can be made to A.G.H.

## Supplementary Information

**Supplementary Figures**

- Supplementary Fig. 1. Enhancers in *MYCN*-expressing neuroblastoma contain binding motifs for core regulatory circuit factors
- Supplementary Fig. 2. Asymmetric amplicon boundaries cannot be explained by experimental platform, clinical variables or chromosomal fragile sites
- Supplementary Fig. 3. H3K27ac ChIP-seq data characteristics on the *MYCN* amplicon for 13 *MYCN*-amplified neuroblastoma cell lines
- Supplementary Fig. 4. Long-read nanopore sequencing enables *de novo* assembly of *MYCN* neighborhoods in neuroblastoma cell lines.
- Supplementary Fig. 5. *De novo* assembly confirms close co-amplification of *MYCN* and e4 in Kelly and NGP
- Supplementary Fig. 6. Enhancer hijacking and neo-TAD formation on the *MYCN* amplicon in IMR-5/75 and CHP-212
- Supplementary Fig. 7. Fluorescence in situ hybridization of *MYCN* ecDNA in CHP-212 and *MYCN* HSRs in IMR-5/75, Kelly and NGP

---

**Supplementary Fig. 1.**
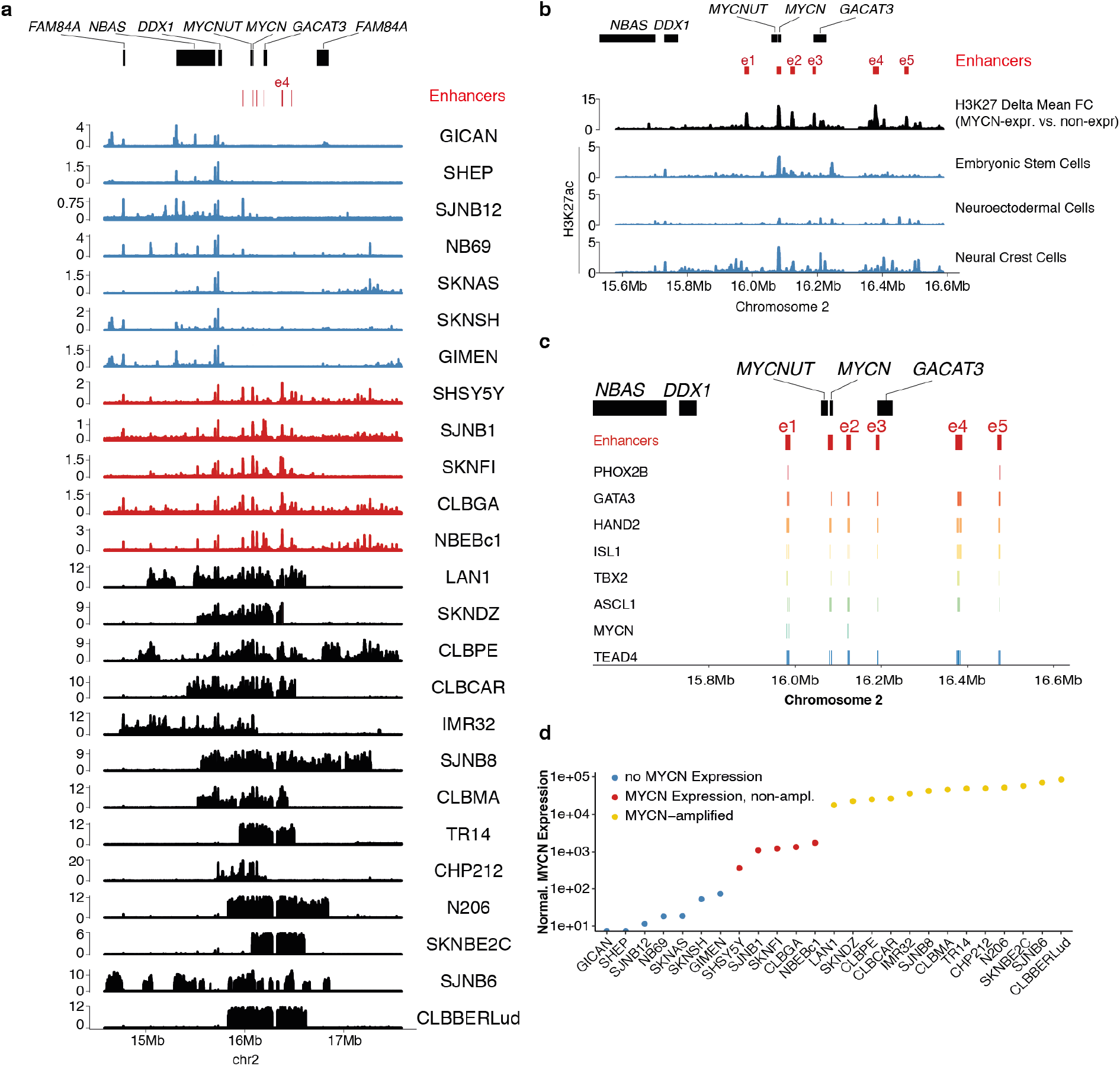
Enhancers in MYCN expressing neuroblastoma contain binding motifs for core regulatory circuit factors. **a** H3K27ac ChIP-seq signal (counts per million in 10bp bins, smoothed in 1kb bins) for seven non-*MYCN*-expressing neuroblastoma cell lines (blue), five *MYCN*-expressing non-*MYCN*-amplified neuroblastoma cell lines (red) and 13 MYCN-amplified neuroblastoma cell lines. **b** Differential composite H3K27ac signal for *MYCN*-non-expressing vs. *MYCN*-expressing non-*MYCN*-amplified cells (difference in the group-wise mean fold change H3K27ac vs. input) and H3K27ac ChIP-seq (counts per million in 10bp bins, smoothed in 1kb bins) signal for *in vitro* differentiated developmental cell types (embryonic stem cells, neuroectodermal cells, neural crest cells). **c** Core regulatory circuit factor (PHOX2B, GATA3, HAND2, ISL1, TBX2, ASCL1), MYCN and TEAD4 binding motif positions in the *MYCN*-driving enhancers e1-e5. Only motif hits within the enhancer regions are depicted. **d** *MYCN* expression for 25 neuroblastoma cell lines as determined by RNA-seq (size factor-normalized read counts) classified into no *MYCN* expression (size factor normalized expression lower than 100), *MYCN*-expressing non-*MYCN*-amplified cells (size factor normalized expression 100 or more) and *MYCN*-amplified cell lines.

**Supplementary Fig. 2.**
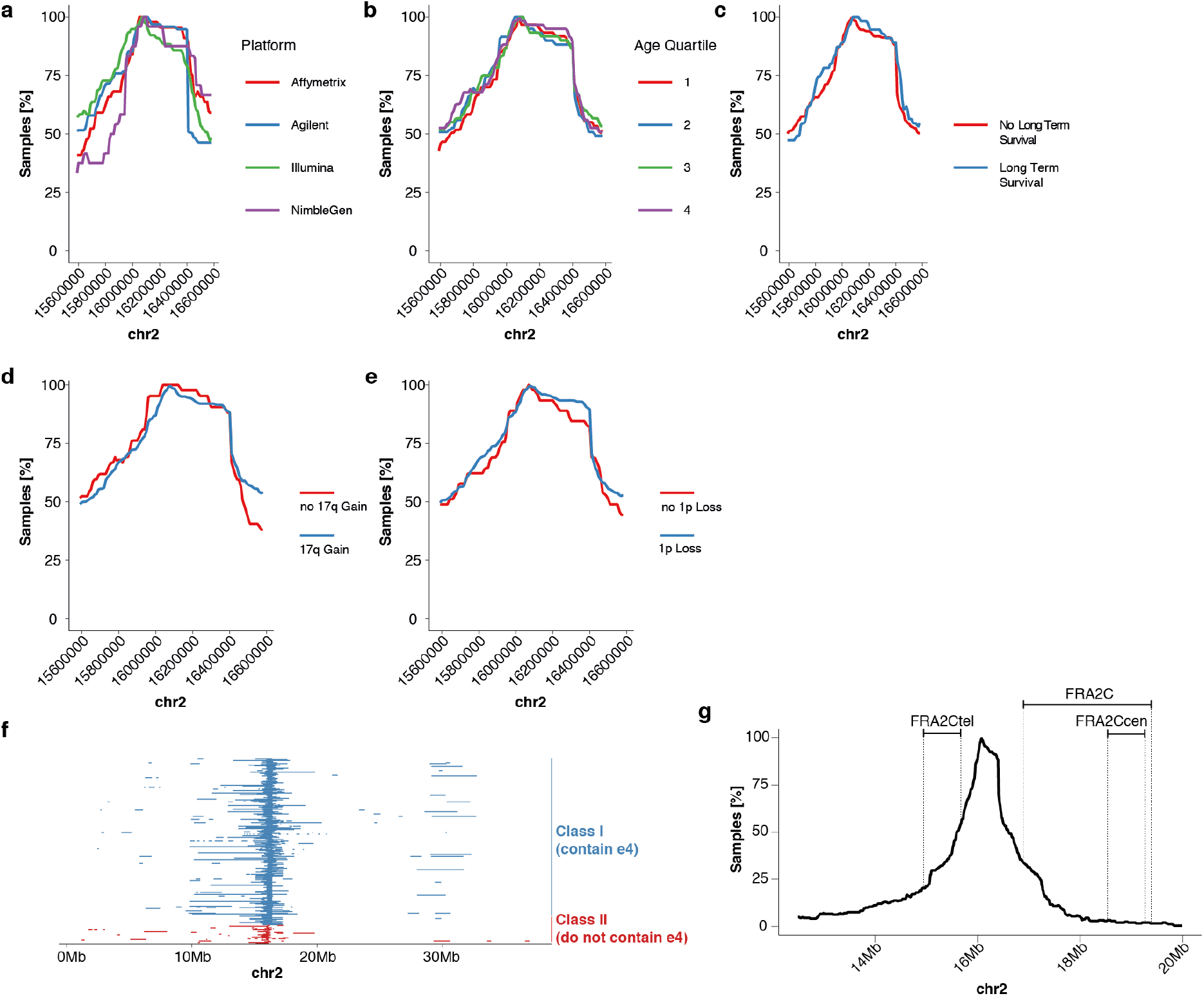
Asymmetric amplicon boundaries cannot be explained by experimental platform, clinical variables or chromosomal fragile sites. **a-e** Percent coamplification in 10kb bins for *MYCN*-amplified neuroblastoma (n=240) split by the experimental method to measure genomic copy-number (**a**, Affymetrix SNP array, Agilent aCGH platform, Illumina SNP array, NimbleGen aCGH platform), the age quartile of patients (**b**, 1=lowest quartile, 4=highest quartile), long-term survival (**c,** defined as survival beyond five years post diagnosis) and the genetic factors 17q gain (**d)** and 1p loss (**e**). **f** Amplified regions on chromosome 2 (0Mb-40Mb) for primary neuroblastoma (n=240), colored by amplicon class (class I amplicons including e4 vs. class II amplicons not including e4) **g** Percent co-amplification in 10kb bins around *MYCN* for *MYCN*-amplified neuroblastoma (n=240) and position of common fragile sites on chromosome 2 between 12Mb and 20Mb.

**Supplementary Fig. 3.**
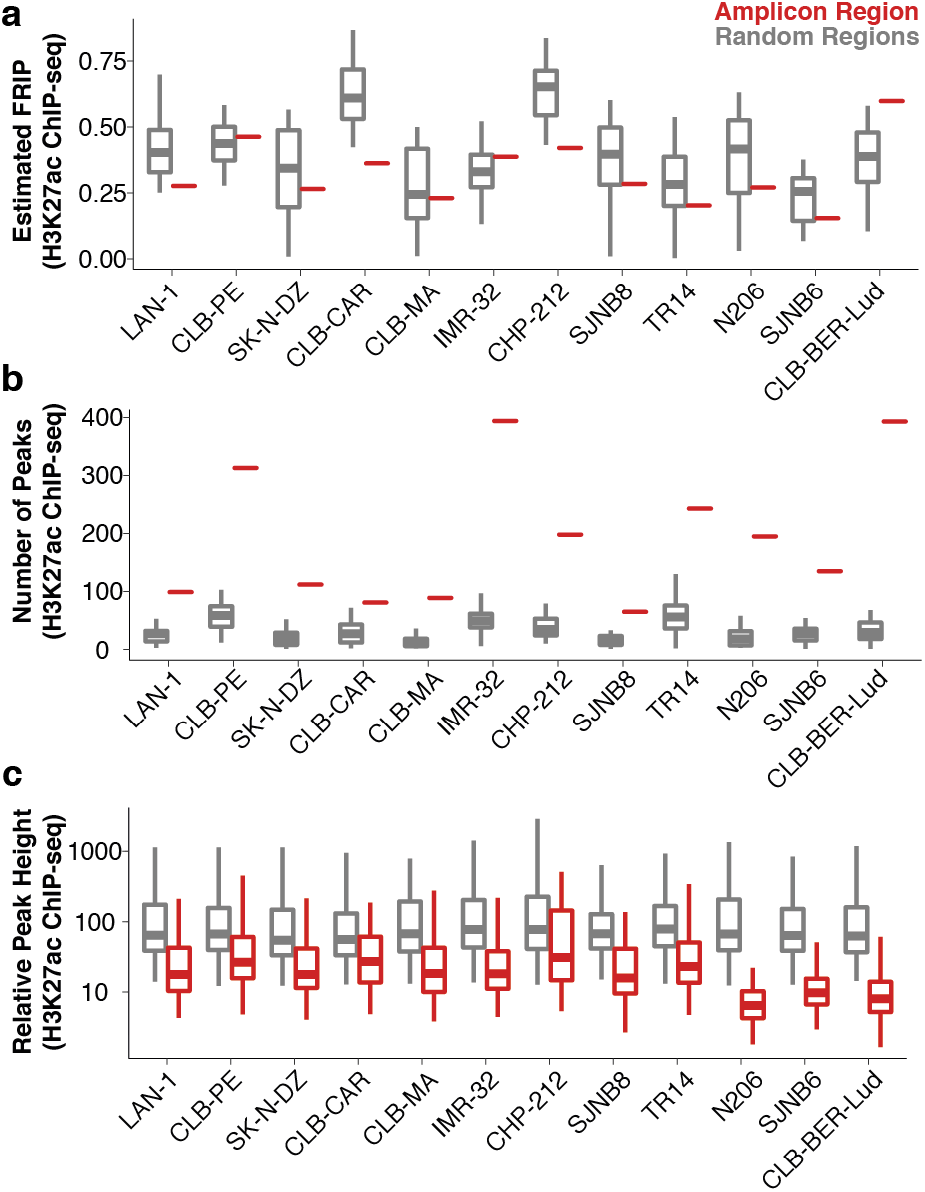
H3K27ac ChIP-seq data characteristics on the *MYCN* amplicon for 13 *MYCN*-amplified neuroblastoma cell lines. **a** Estimated fraction of H3K27ac reads in peaks for amplified regions (red) vs. randomly drawn genomic regions of matching size (grey, n=30) in 12 *MYCN*-amplified neuroblastoma cell lines. **b** Number of peaks for amplified regions (red) vs. randomly drawn genomic regions of matching size (grey, n=30) in 12 *MYCN*-amplified neuroblastoma cell lines. **c** Relative H3K27ac peak heights (compared to amplicon background) for amplified regions (red) vs. randomly drawn genomic regions of matching size (grey, n=30). In all boxplots, boxes depict the median, the upper quartile boundary and the lower quartile boundary. Whiskers extends to the largest data point within 1.5-fold of the interquartile range.

**Supplementary Fig. 4.**
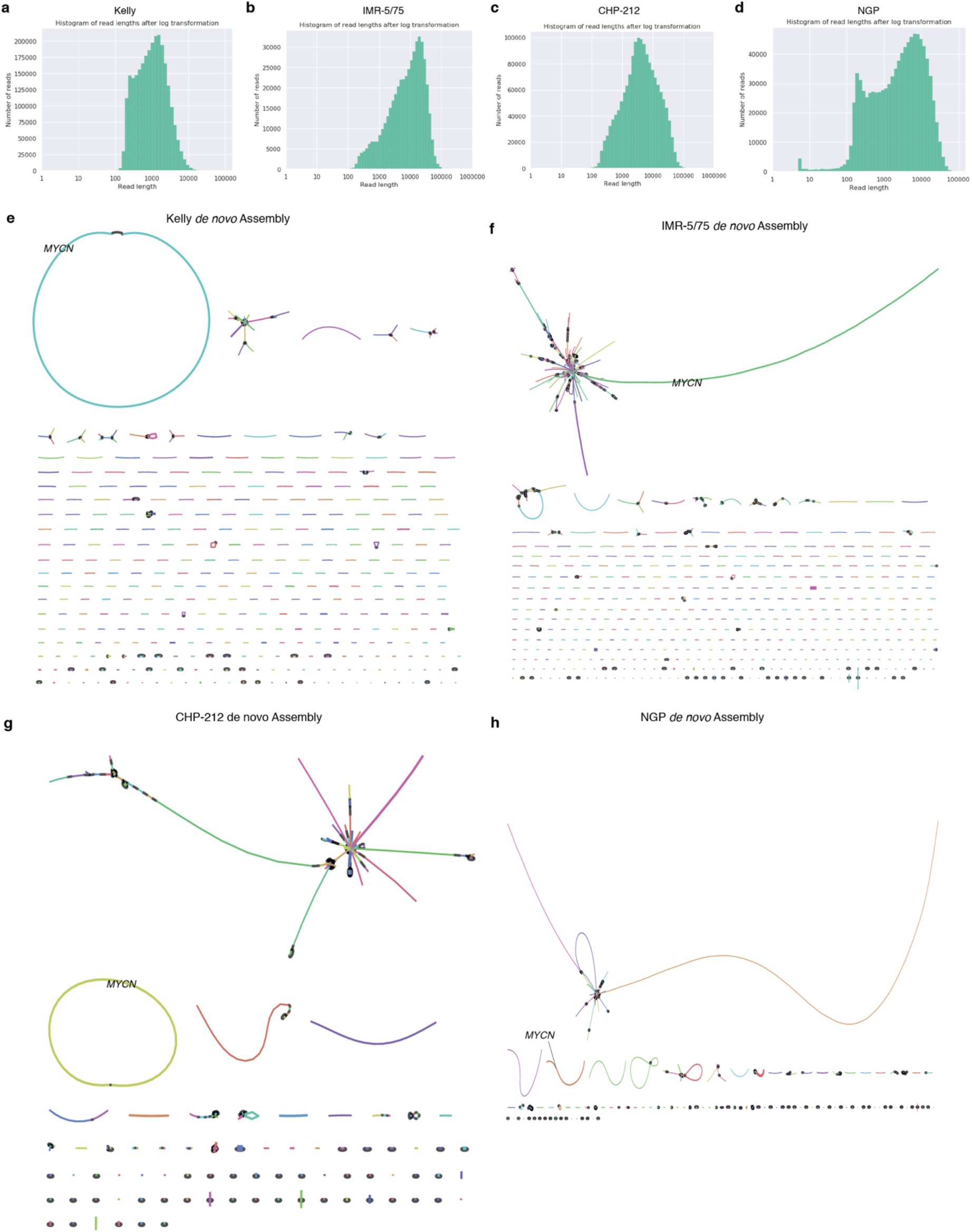
Long-read nanopore sequencing enables *de novo* assembly of *MYCN* neighborhoods in neuroblastoma cell lines. **a-d** Nanopore read length distribution (log-transformed) for the neuroblastoma cell lines Kelly (**a**, n=2,654,406), IMR-5/75 (**b**, n=474,980), CHP-212 (**c**, n=1,554,048) and NGP (**d**, n=952,031). **e** Nanopore long read-based *de novo* assembly of Kelly cells yields 477 contigs and an overall assembly N50 of 24,929 bp. BLAST analysis locates *MYCN* on a circular 975,932 bp contig. **f** Nanopore long read-based *de novo* assembly of IMR-5/75 cells yields 6,265 contigs and an overall assembly N50 of 91,273 bp. BLAST analysis locates *MYCN* on a linear 3,201,197 bp contig. **g** Nanopore long read-based *de novo* assembly of CHP-212 cells yields 21,264 contigs and an overall assembly N50 of 113,845 bp. BLAST analysis locates *MYCN* on a circular 1,705,218 bp contig. **h** Nanopore long read-based *de novo* assembly of NGP cells yields 6,550 contigs and an overall assembly N50 of 60,981 bp. BLAST analysis locates *MYCN* on a linear 623,907 bp contig.

**Supplementary Fig. 5.**
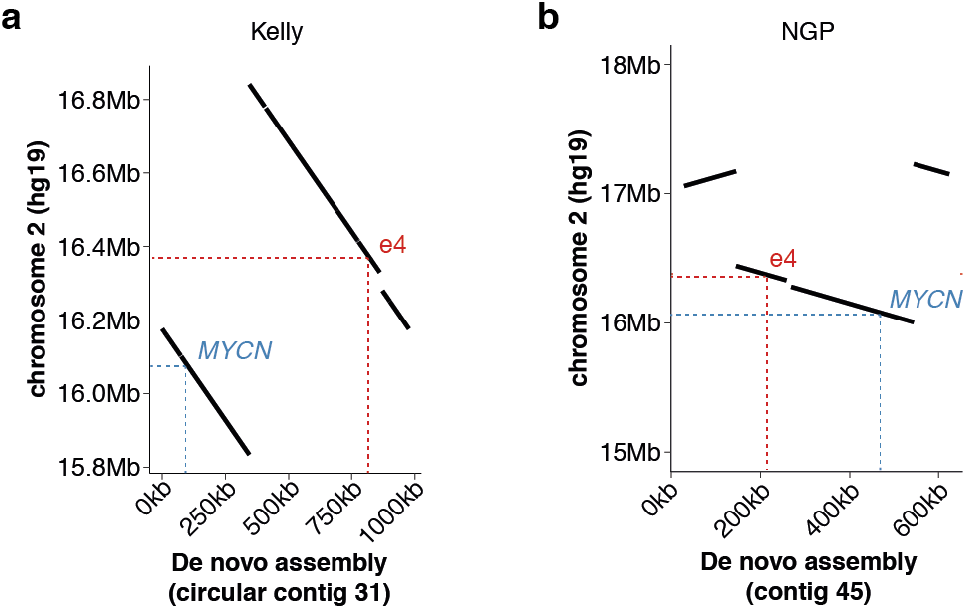
*De novo* assembly confirms close co-amplification of *MYCN* and e4 in Kelly and NGP. Mapping of the *de novo* assembled *MYCN*-containing contig to hg19 in Kelly (**a**) and NGP (**b**) cells. Positions of *MYCN* and e4 are marked on the contig and in the reference genome. Note that the Kelly contig is circular such that the shortest distance from e4 to MYCN spans the contig circle junction.

**Supplementary Fig. 6.**
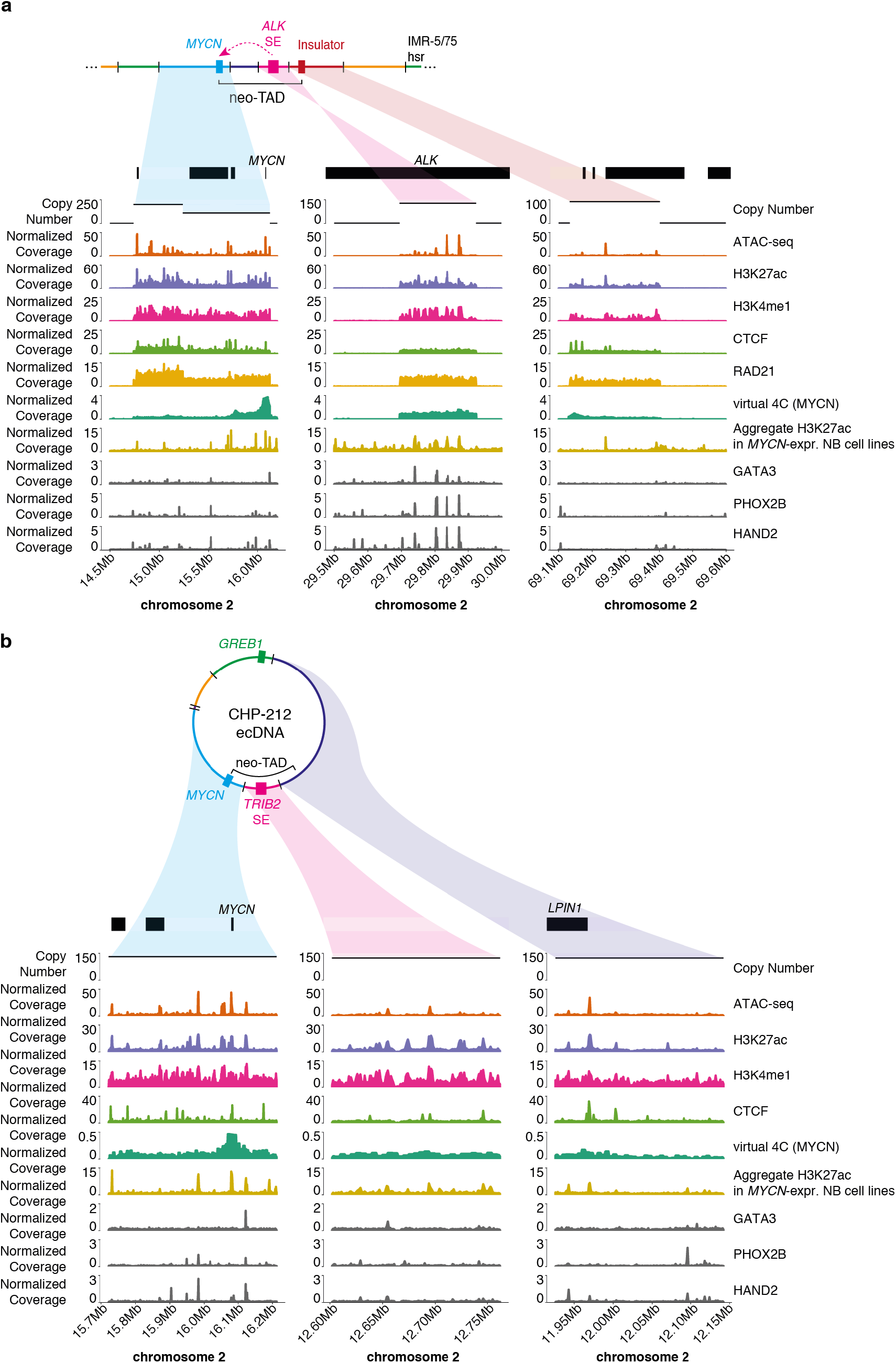
Enhancer hijacking and neo-TAD formation on the *MYCN* amplicon in IMR-5/75 and CHP-212. Regions that contribute to neo-TAD formation are depicted for IMR-5/75 (**a**) and CHP-212 (**b**). Top to bottom: Gene bodies, Copy Number, ATAC-seq, H3K27ac ChIP-seq, H3K4me1 ChIP-seq, CTCF ChIP-seq, RAD21 ChIP-seq (only available for IMR-5/75), virtual 4C with *MYCN* as the viewpoint (mean Knight-Ruiz normalized interaction frequency of three 5kb bins (chr2:16,075,000-16,085,000) around *MYCN*), the aggregate H3K27ac signal over 7 *MYCN*-expressing non-MYCN-amplified neuroblastoma celllines (mean fold change over input), GATA3 ChIP-seq, PHOX2B ChIP-seq and HAND2 ChIP-seq (all CRC transcription factors in the neuroblastoma cell lines CLB-GA). ChIP-seq and ATAC-seq is depicted as counts per million in 10bp bins, smoothed in 1kb bins.

**Supplementary Fig. 7.**
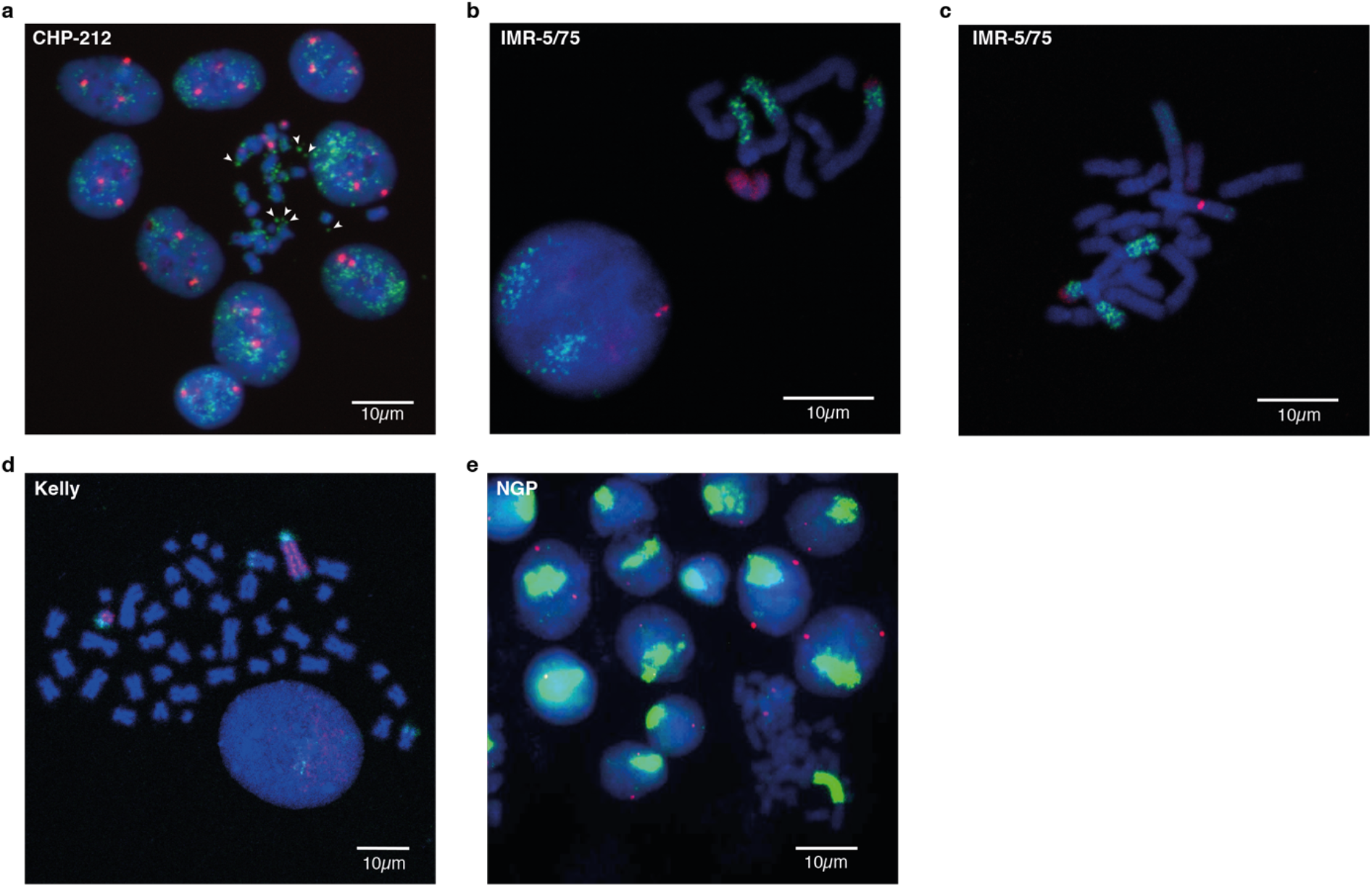
Fluorescence in situ hybridization of *MYCN* ecDNA in CHP-212 and *MYCN* HSRs in IMR-5/75, Kelly and NGP. **a** Fluorescence in situ hybridization (FISH) of CHP-212 metaphase spreads with a *MYCN* probe (green) and a probe for the chromosome 2 centromere (red). Arrowheads point to *MYCN* ecDNA. **b, c** FISH of metaphase spreads in IMR-5/75 with a *MYCN* probe (green), a chromosome 2 centromere (red) and a chromosome 12 paint (red) **d** FISH of metaphase spreads in Kelly cells with a *MYCN* probe (red) and chromosome 17 paint (green). **e** FISH of NGP metaphase spreads with a *MYCN* probe (green) and a probe for the chromosome 2 centromere (red).

